# CRISPR-Cas9 Gene Editing in Lizards Through Microinjection of Unfertilized Oocytes

**DOI:** 10.1101/591446

**Authors:** Ashley M. Rasys, Sungdae Park, Rebecca E. Ball, Aaron J. Alcala, James D. Lauderdale, Douglas B. Menke

**Author notes:** Correspondence: Douglas B Menke, University of Georgia, Department of Genetics, 500 DW Brooks Dr, Athens, GA 30602, USA.

## Abstract

CRISPR-Cas9 mediated gene editing has enabled the direct manipulation of gene function in many species. However, the reproductive biology of reptiles presents unique barriers for the use of this technology, and there are currently no reptiles with effective methods for targeted mutagenesis. Here we present a new approach that enables the efficient production of CRISPR-Cas9 induced mutations in *Anolis* lizards, an important model for studies of reptile evolution and development.

## Main

Squamates (lizards and snakes) comprise a diverse group of reptiles represented by over 10,000 recognized species^1^. However, mechanistic studies of gene function in squamates and other reptiles lag behind other major vertebrate groups. While the adoption of CRISPR-Cas9 mediated gene editing has enabled direct manipulation of gene function in many fish^2,3^, amphibian^4,5^, avian^6^, and mammalian species^7,8^, there remain no reptilian model systems with established methods for the production of targeted sequence alterations. To date, attempts to manipulate gene function in reptiles have been limited to a small number of studies employing whole embryo culture coupled to viral-or electroporation-based methods to alter gene expression^9,10^. These methods produce transient, localized, and highly mosaic patterns of transgenesis. Moreover, these techniques have not been used to engineer targeted gene modifications in any reptile species.

Among squamates, *Anolis* lizards are compelling candidates for the establishment of gene editing methods. Over the past 50 years, anoles have become one of the central model systems for studies of reptile evolution, physiology, and development^11^. This group has experienced an extensive adaptive radiation in the Caribbean with hundreds of described species that display a wide range of morphological, behavioral, and physiological differences. Studies of the convergent evolution of similar sets of *Anolis* “ecomorphs”, or habitat specialists, on different Caribbean Islands has produced a rich-literature on the biology of *Anolis* lizards. Although many *Anolis* species can be successfully raised in the lab, we chose to develop genome editing in the brown anole lizard, *Anolis sagrei*. This invasive lizard is now found far beyond its native Caribbean range and is ideal for genetic studies due to its small size, ease of husbandry, long breeding season, and relatively short generation time.

CRISPR-Cas9 mediated genome editing is an effective method for producing genetically modified vertebrates^12^. Typically, CRISPR-Cas9 components are microinjected into vertebrate embryos at the one-cell stage to generate individuals potentially harboring alterations at the locus of interest. However, there are significant challenges associated with microinjection of *Anolis* zygotes. These challenges include internal fertilization and the long-term storage of sperm within the oviducts of adult females, which makes timing the microinjection of single cell embryos extremely difficult. At the time of ovulation *Anolis* eggs are also quite large (∼8 mm in length) and are filled with substantial amounts of yolk; these oocytes are fragile and are difficult to manipulate without rupturing. Furthermore, after fertilization the egg shell must be deposited around the egg and embryonic development is initiated before the egg is laid. Finally, unlike the hard shells of birds, the egg shells that enclose *Anolis* embryos are pliable and no air space is present within the egg, presenting obstacles for embryo manipulation within the egg shell. Most of these reproductive challenges for microinjection are not unique to anoles, but are features that are typical of many reptiles. Here we demonstrate the use of CRISPR-Cas9 to mutate the *tyrosinase* gene in *Anolis sagrei* using an approach that is generalizable to other loci. The procedure we have developed involves microinjection of CRISPR-Cas9 components into immature, unfertilized *Anolis* oocytes within the ovaries of female lizards.

Reproductively active *A. sagrei* females lay approximately 1 egg every week, similar to *Anolis carolinensis*^13^. Each ovary contains a series of approximately 10 maturing ovarian follicles arranged by size, with the smallest follicle closest to the germinal bed and the largest vitellogenic follicle positioned distally (Fig. S1). With the exception of the largest follicle, these follicles are previtellogenic or in the early stages of vitellogenesis and are largely similar in size between the left and right ovaries. In our approach, female lizards are anesthetized and are placed on a surgical platform underneath a standard dissecting scope. Left and right ovaries are separately accessed via vertical incisions positioned along the left or right flank, respectively (Fig. 1). During surgery, the ovary can be gently moved to allow easy observation and injection of the oocytes under a dissecting microscope. Oocytes are microinjected with Cas9 ribonucleoprotein complex (Cas9 RNP) while remaining associated with the ovary (Fig. 1, Fig. S1, and Video S1). In our hands, oocytes that are 0.75 to 5 mm in diameter yield the highest frequency of mutant animals (Fig. S2). A typical anole ovary has 4 to 6 oocytes in this range. Therefore, approximately 10 oocytes in this size range can be injected per animal. Oocytes equal to or larger than 6 mm in diameter are not injected due to the increased risk of rupturing these large, yolk-filled oocytes. After microinjection of oocytes is completed on one side, the incision is sealed with veterinary glue. The procedure is then repeated on the opposite side. We routinely inject 5 females per day for a total of about 50 oocytes per injection session.

**Figure 1.**
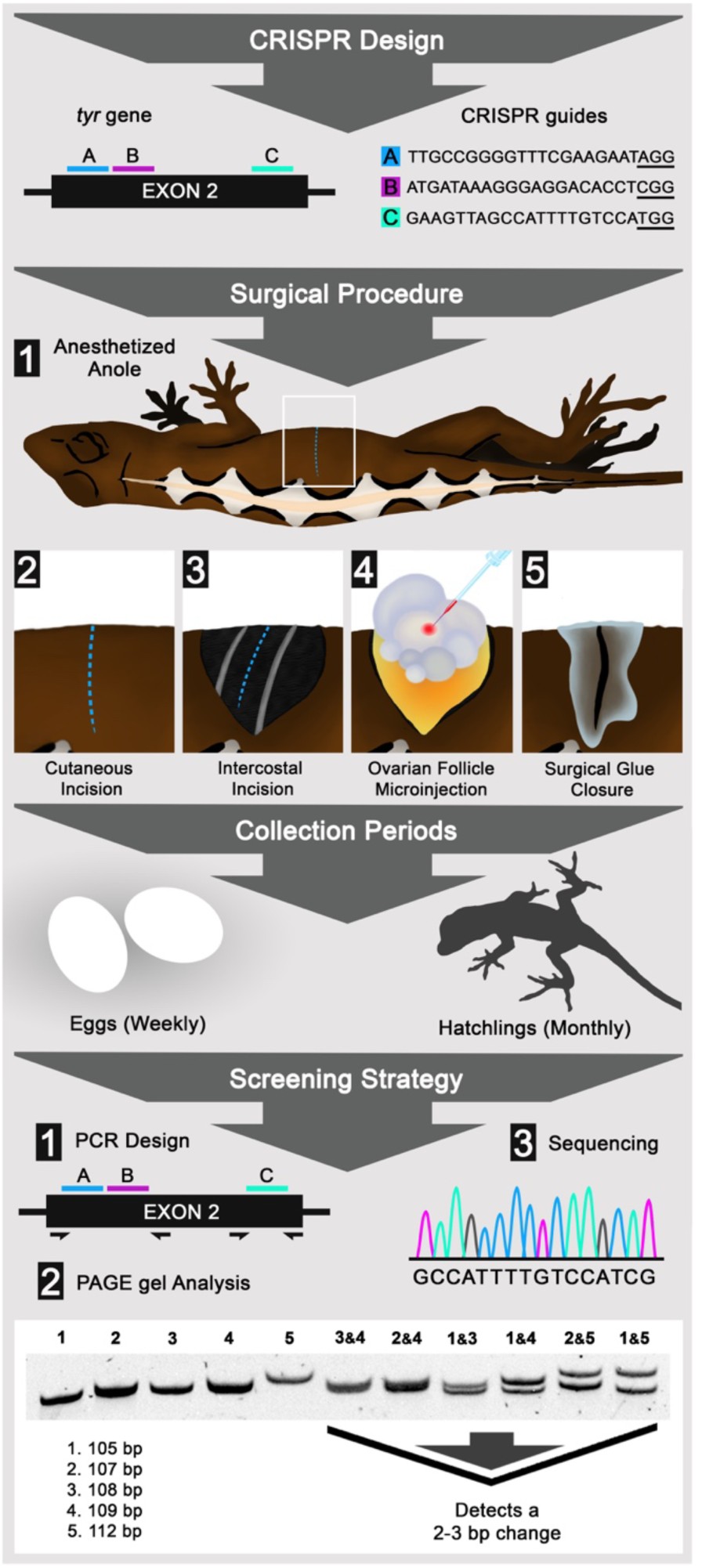
Gene editing in lizards through microinjection of ovarian follicles. Flow diagram detailing CRISPR design, surgical procedure, collection periods, and screening strategy. CRISPR design shows the placement and sequence of CRISPR guides A (blue), B (pink), and C (cyan) within exon 2 of the *tyr* gene; PAM sites are underlined. The surgical procedure panel depicts lizard anesthesia and surgical steps to access and microinject ovary follicles. The collection periods panel highlights the time between gathering eggs and raising hatchlings. The screening strategy panel illustrates the steps used to detect *tyr*-crispants including: 1) PCR primer design, 2) PAGE analysis, which can reliably detect 2-3 bp changes, and 3) Sanger sequencing.

Following recovery, the microinjected oocytes continue to mature within the female and are eventually ovulated and fertilized through natural mating with an introduced male or via stored sperm from previous matings. The ovaries have an ovulatory cycle that ranges from two to four weeks, but the left and right ovaries are out of phase relative to each other, which results in eggs laid in alternating order from follicles on the left and right ovaries^14^. Because of the alternating follicle contributions from the left and right ovaries, a putative “follicle train” can be established by ranking follicles based on size and by designating ovarian turns. By ordering follicles in this manner, one can infer the range of follicle sizes which yielded crispant lizards. For instance, follicles in the 3 to 5 mm range develop into eggs that are laid within three to four weeks post injection (Fig. S2).

To assess the effectiveness of our approach, we targeted the second exon of the *tyrosinase* (*tyr*) gene. *Tyrosinase* was chosen for this study because loss-of-function mutations are viable in a wide-range of vertebrates, the resulting pigmentation phenotypes are readily detected, and this allowed us to develop a new *Anolis* model to investigate human albinism. Cas9 protein coupled to a mixture of three different synthetic *tyr* guide RNAs was injected into immature oocytes. The decision to simultaneously inject three guide RNAs was motivated by the presence of a number of single-nucleotide polymorphisms (SNPs) in *tyr* exon 2 within the population of lizards used for these experiments. For these experiments, a total of 146 oocytes from 21 reproductively active females were microinjected over the course of eight surgical sessions.

We obtained nine F0 animals harboring mutations in *tyr* exon 2. Four of these animals were phenotypically albino and harbored loss-of-function mutations at both *tyr* alleles. Five animals carried heterozygous loss-of-function mutations and exhibited normal pigmentation. Mutant alleles could be visualized by PAGE after PCR amplification across *tyr* exon 2 using genomic DNA prepared from F0 embryos or hatchlings as a template. The changes associated with each of the CRISPR-Cas9 induced mutations was determined by sequencing the amplicons from each animal and comparing the sequence to that of wild-type lizards in the colony. The overall mutation frequency in terms of mutant lizards per follicle injected was 6.2%. A mutation frequency of 9.7% was obtained from microinjected follicles that were 1.5 to 2.5 mm in diameter, while follicles 0.75 to 1.0 mm and 3 to 5 mm in diameter yielding frequencies of 9.3% and 5.6%, respectively. No mutations were obtained from microinjection into follicles smaller than 0.5 mm in diameter. Consistent with results in other vertebrates, CRISPR-Cas9 genome editing in lizards resulted in indels that typically ranged in size from 3 to 17 base pairs.

Our discovery that microinjection of genome editing reagents into oocytes generates F0 lizards carrying bi-allelic mutations in the targeted gene, demonstrates that our approach offers an efficient path for directly testing gene requirement in anoles. For genes where loss of function mutations are homozygous lethal, or studies requiring heterozygous animals, we expect that naturally occurring SNPs can be used to facilitate targeting of one allele only.

The establishment of CRISPR-Cas9 editing in this inexpensive reptilian system will finally permit mechanistic studies of gene function to be performed in reptiles. We anticipate the gene-editing strategy we have developed in anoles can also be successfully applied to many other squamate species.

## Acknowledgments

We thank S. Divers for his guidance on reptile anesthesia and surgical techniques and J. Eggenschwiler and C. Sabin for comments on this manuscript. This work was funded by National Science Foundation awards #1149453 to D.B.M. and #1827647 to D.B.M. and J.D.L., and a Society for Developmental Biology Emerging models grant to A.M.R. A.M.R. was supported by NIH training grant T32GM007103 and by an ARCS Foundation Scholarship.

## Author contributions

A.M.R., J.D.L., and D.B.M conceived and designed the experiments. A.M.R., performed the majority of the experiments with support from S.P., A.J.A. and R.E.B. Data interpretation was carried by A.M.R., S.P., R.E.B., J.D.L., and D.B.M. The manuscript was written by A.M.R., D.B.M., and J.D.L. with help from all other authors.

## Competing interests

The authors declare no competing interests.

**Figure S1.**
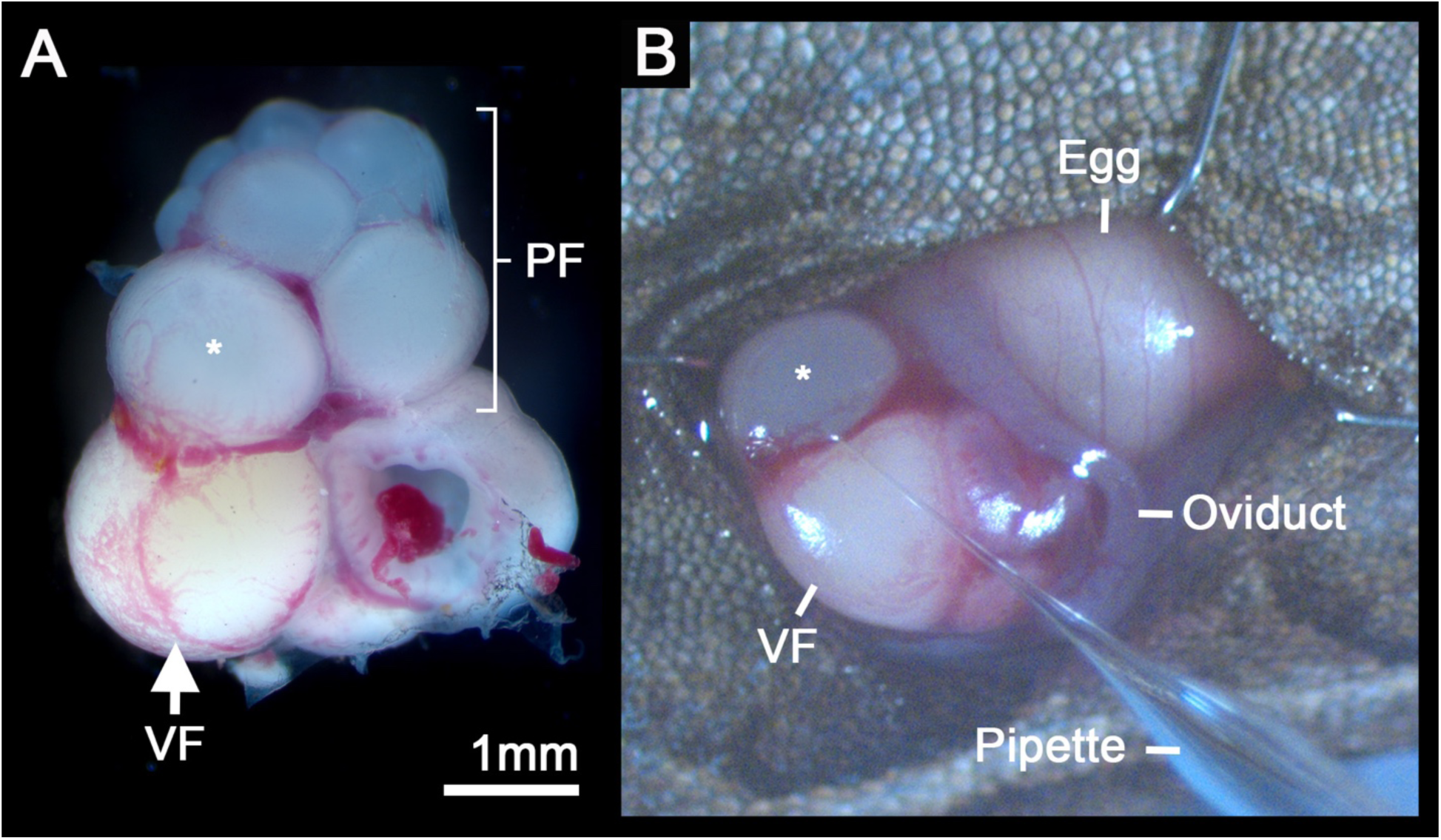
Anole lizard ovary. (A) Dissected ovary showing previtellogenic (PF) and vitellogenic (VF) follicles. (B) Same ovary prior to dissection showing microinjection of a 1.5 mm diameter follicle (asterisk in panels A and B).

**Figure S2.**
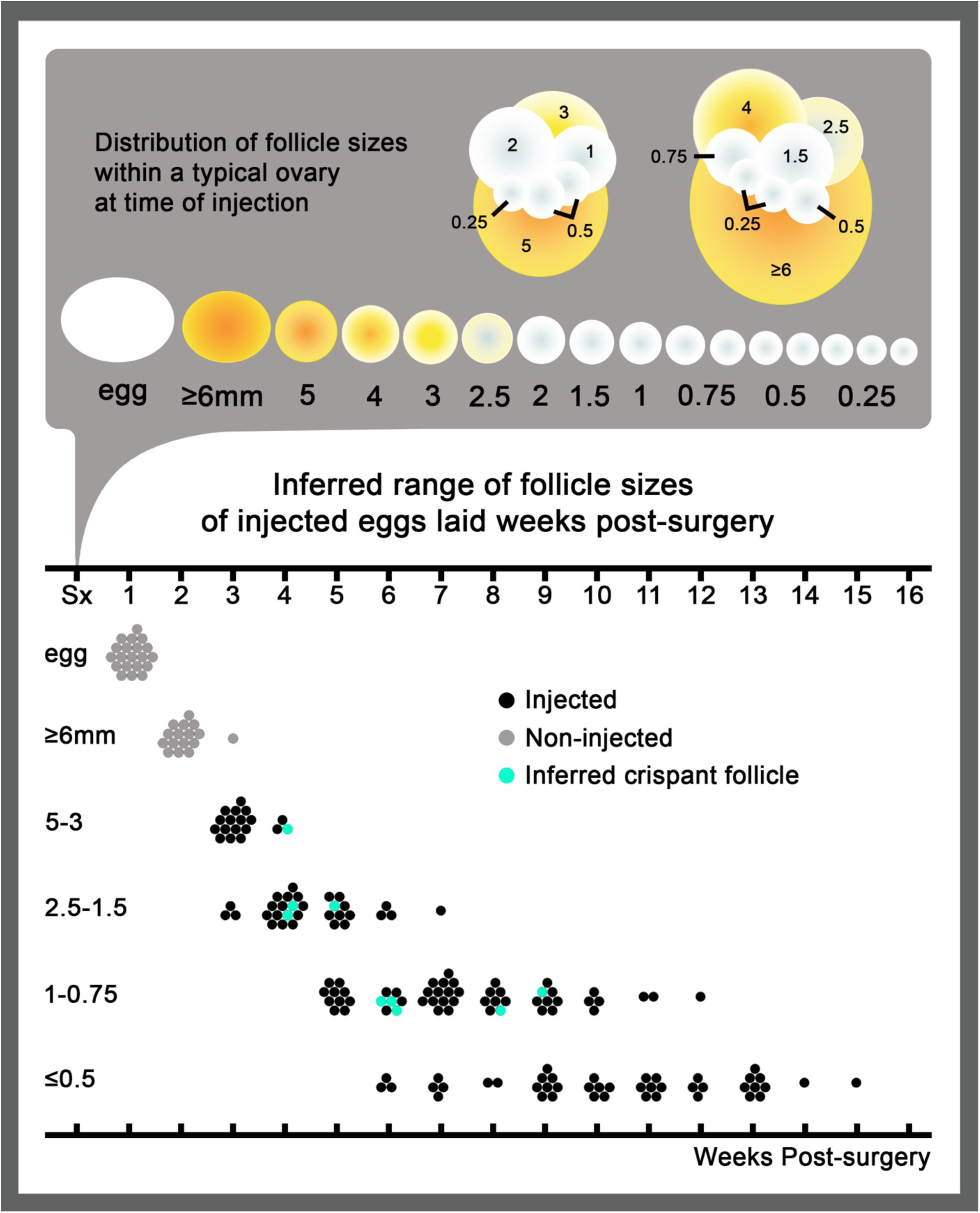
Inferred relationship between follicle size and crispants. (Top) the distribution of follicles sizes within ovaries at time of injection depicted as a “follicle train.” (Below) a distribution graph showing inferred follicle sizes over weeks post-surgery. Non-injected eggs and large yolky follicles are shown in grey, while injected follicles are in black and follicles that likely produced crispant lizards are in cyan.

## Methods

### Animals

Animals used in this study were wild-caught *Anolis sagrei* from Orlando, FL. Lizards were housed at University of Georgia following published guidelines^15^. Breeding cages housed up to 4 females and 1 male together. Twenty-one adult females from cages that consistently produced eggs were selected for this study. All experiments followed the National Research Council’s Guide for the Care and Use of Laboratory Animals and were performed with the approval and oversight of the University of Georgia Institutional Animal Care and Use Committee (A2016 09-008-Y2-A3).

### Selection of crRNA guide sequences and preparation of Cas9 RNP

The CRISPOR target selection tool (version 4.4) was used to select target regions with efficiency scores of 50% or greater within the second exon of the *A. sagrei tyr* gene^16^. The *tyr* gene reference sequence was obtained from a draft genome assembly of *Anolis sagrei*. Alt-R CRISPR-Cas9 crRNAs, tracrRNA, and Cas9 V3 were purchased from Integrated DNA Technologies, Inc. The crRNA target sequences were as follows:

*AsagTyrEx2A*: 5′-TTGCCGGGGTTTCGAAGAAT-3′

*AsagTyrEx2B*: 5′-ATGATAAAGGGAGGACACCT-3′

*AsagTyrEx2C*: 5′-GAAGTTAGCCATTTTGTCCA-3′.

Cas9 RNP was prepared by following manufacturer recommendations. A 5 µM injection solution was made using standard microinjection solution (10 mM Tris-HCl, pH 7.4) containing phenol red to help verify that injected solutions entered the oocytes.

### Anesthesia and analgesia

Lizards were anesthetized by administering 30 mg/kg of Alfaxalone (Alfaxan, 10 mg/mL, Jurox) in combination with 0.1 mg/kg of dexmedetomidine (Dexdomitor 5 mg/10 mL, Zoetis/Orion). To ensure accurate dosing, these drugs were administered by subcutaneous injection in the cervical area as an Alfaxalone/Dexmedetomidine (A/D) mixture. Preoperative analgesia was obtained by subcutaneous injection of meloxicam (0.3 mg/kg, Loxiject, 5 mg/mL, Henry Schein) in the dorsal epaxial area just above the shoulder and topical application of lidocaine (2.0 mg/kg, Lidocaine HCL, 2%, Hospira) over the surgical site. See video for injection sites. The anesthetic combination A/D provided approximately 30 min of surgical anesthesia time. Lizards typically recovered about 40-45 min following A/D administration. If a longer anesthesia time was required, a second dose of 30 mg/kg of alfaxalone alone was administered 25-30 min post induction dose, providing an additional 30 min of surgical anesthesia.

The method and location where injections were made were specifically chosen to avoid some of the challenges with administering drugs to reptiles. One such issue to be aware of is the hepatic-first pass effect which is a phenomenon found in many reptiles where, drugs, if administered in hindlimb or caudal regions, are rapidly cleared by the ventral abdominal and hepatic portal veins and metabolized by the liver, inhibiting wide systemic circulation. We have found that administering A/D subcutaneously in the dorsal epaxial area just above the shoulder region resulted in only moderate to light levels of anesthesia. Contrary to this, lizards administered subcutaneously in cervical area are rapidly induced and reach a surgical plane of anesthesia within 1 min. Because injection volumes can be large, a subcutaneous method was also preferred over an intramuscular approach. For these reasons, the cervical area and a subcutaneous route were used in this study.

Another important factor that can influence drug metabolism in reptiles is body temperature. Lizards are ectothermic and are dependent upon the environmental temps to regulate their body warmth which in turn impacts their metabolic rate. A decrease in body temperature will lead to a decrease in drug metabolism potentially resulting in a persistence of circulating drugs. In such an instance, this can result in an animal responding poorly, prolongment of anesthesia recovery time, or in some cases lead to death. The converse is also true. Animals maintained at too high a temperature may metabolize and clear drugs quickly resulting in insufficient anesthesia time. To avoid this, anesthetized anoles should be maintained on a heating source at around 32°C until fully recovered.

### Surgery and microinjection

After successful anesthesia induction, the lizard becomes non-responsive to any noxious stimuli (i.e., an absence of response to a cloacal/tail clamp that normally induces severe discomfort). The anesthetized lizard was placed into right lateral recumbency and the left flank was aseptically prepared by alternating disinfection with 70% ethanol and 7.5% povidone-iodine (Surgical Scrub Solution, 16 fl. oz. 473 mL, Dynarex) wipes for 5 minutes.

Following standard surgical practices, sterile iris scissors (FST, item 15023-10) were used to make an 8-10 mm vertical cutaneous incision on the left side, in the mid-coelom region. A second incision between the ribs was made through the musculature (i.e., internal/external intercostal and pigmented coelom muscle layers) to enter the coelom. The ovary can be found dorsally in the mid-coelom region and was easily accessible by shifting intestines gently aside using blunt forceps (FST: 45° angled forceps, item 00649-11; FST: strait forceps, item 00632-11). Once located, the ovary was carefully rotated and repositioned to expose immature follicles ranging anywhere from 0.25 mm to 5 mm in size.

Using the blunt forceps to clasp and hold the ovary in place, a microinjection needle was visually guided into the follicle center at an angle between 35-45° degrees relative to horizontal. 5 µM Cas9 RNP solution was then injected into follicles at differing volumes ranging from as little as 15 nL to as much as 575 nL which was dependent upon needle and follicle size. Retrospectively, ideal injection volumes were determined (3≤5 mm, 300-500 nL; 2≤2.5 mm, 200-250 nL; 1≤1.5 mm, 100-150 nL; and 0.75 mm, 25 nL) based on surgical sessions that produced mutants. Large yolky follicles greater than 5 mm in diameter, and eggs already present in the oviduct, were not injected. Sterile drops of P-Lytes solution (Veterinary Plasma-Lyte A Injection pH 7.4) were applied directly on the ovary or in coeliotomy opening throughout the procedure to prevent tissue dehydration.

After injection, the ovary was gently returned into the coelom and overlying musculature and skin was lightly pulled together to close the cavity. Tissue adhesive (3M Vetbond, #1469SB) was carefully applied only to only the external surface of the skin, avoiding the underlying musculature. Once the tissue adhesive was dry, the lizard was re-positioned into left lateral recumbency and the procedure was repeated for a right coeliotomy.

During recovery, triple antibiotic ointment (Bacitracin Zinc, Neomycin Sulfate, Polymyxin B Sulfate) was applied topically to the surgical wounds. Lizards were monitored daily for 1 week for any signs of infection, pain, or inflammation. After recovery from anesthesia, females were housed together with their previous female mates and allowed to recover for 7 days prior to reintroducing the male.

All surgeries were performed using sterilized equipment and tools (i.e. forceps and iris scissors) under a dissecting scope (Zeiss Stemi SV11) with a top light (AmScope 80-LED illuminator). Body temperature was maintained throughout the procedure by placing lizards on a heating platform (Fisher Scientific: model 77, serial # 802N0041CAT 12-594) with surgical towels draped between the heat source and the lizards to provide a barrier. The contact surface temperature was held at 32°C and readings continuously taken using thermometer strips. Each laparotomy was performed within 10-12 minutes. Follicle injections were carried out using a standard zebrafish/xenopus microinjection rig (Harvard Apparatus PLI-100 Pico-Injector) set at 20 PSI with an injection time of 50-60 msecs. Initially, a manual micromanipulator (Märzhäuser Wetzlar; MMJ-rechts: 00-42-107-0000) was used to perform steady needle injections. However, use of this limited the degrees of freedom to inject the ovary from multiple directions and angles, therefore, a simple hand-guided technique using no micromanipulator was ultimately preferred and proved to be more efficient. Injection needles with a gradual taper typically used for zebrafish microinjection were made following the Sutter Instrument Company Pipette Cookbook guidelines using a Flaming/Brown Micropipette Puller (Model P-97) and cut to have a 20 to 40 µm diameter opening.

### Mutation screening

Cages were monitored for a specified number of weeks following surgery which was based on the highest number of follicles injected in an ovary (e.g., if 8 and 5 follicles were injected in the right and left ovaries of one lizard, respectively, a cage housing this lizard would be monitored for n=16 weeks). Because these females often had 1 or 2 eggs in the oviduct as well as 2 large (>5mm) un-injected large follicles, cages were monitored for an additional 3-4 weeks following surgery.

Embryos and hatchlings from surgery cages were screened via PCR PAGE analysis under conditions that reliably detect a 2-3bp change^17^. DNA was extracted from tail clips from hatchlings or from tissue collected from embryos following standard protocols. PCR was performed using the following primers: P1, 5-CAAGAACTTTGCAATGGAACAAATG-3’; P2, 5’-GAATTCAACGTCTGCTGAAGATG-3’; P3, 5’-TGTTTAAGTCTGACTCAGTACGAAG-3’; P4, 5’-GGATTACCTTCCAAAGTATTCCTG-3’. See Figure 2 for primer location relative to targeting sites. Sanger sequencing was performed on PCR products to fully characterize mutations.

**Figure 2.**
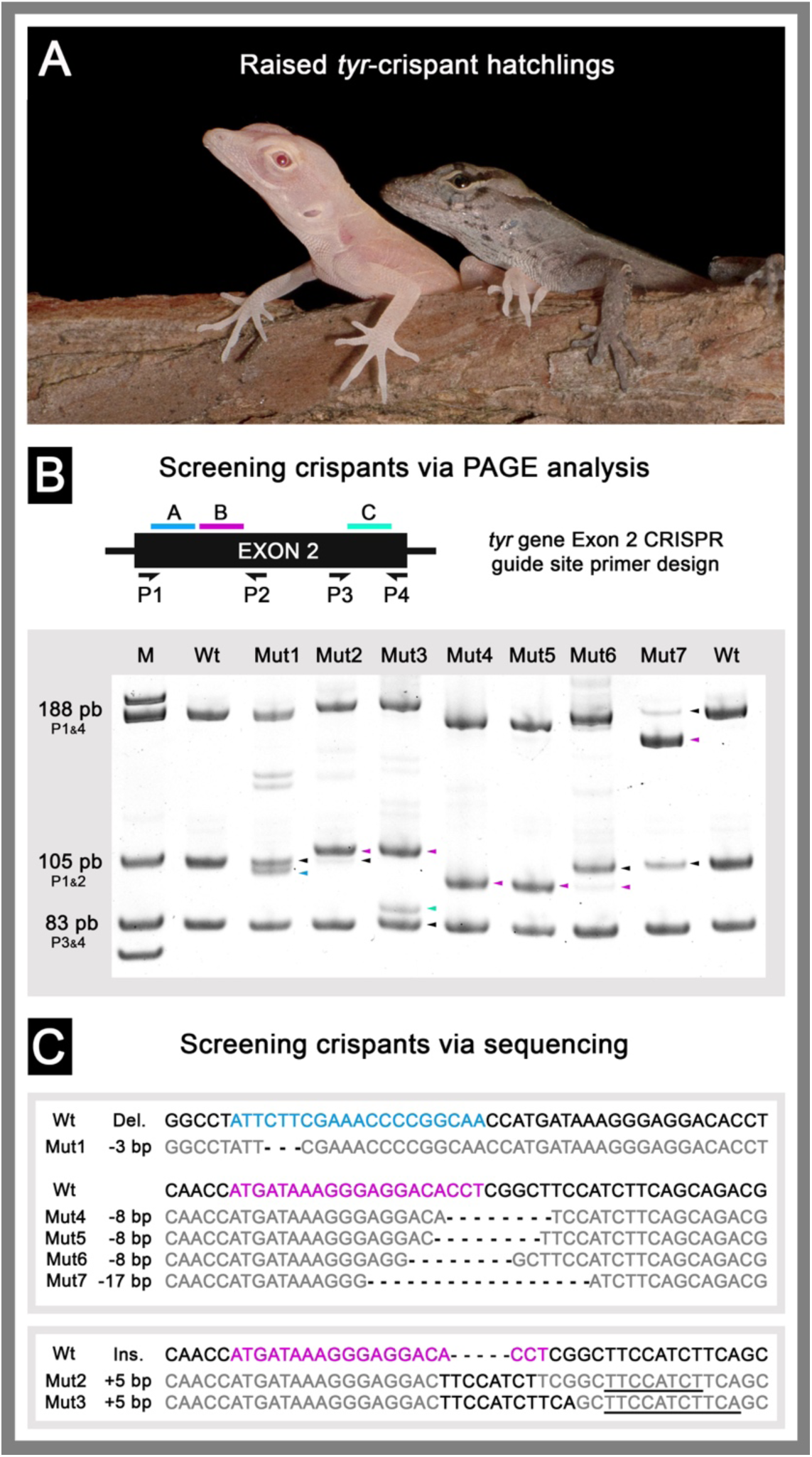
Detection and sequencing of *tyr*-crispants. (A) albino *tyr-*crispant (left) and wildtype (right) aged-matched hatchlings. (B) PCR primer placement (P1-P4) relative to CRISPR target sites A (blue), B (pink), and C (cyan). Representative PAGE results are shown for seven of the mutant lizards. Colored arrows denote bands with altered mobility relative to wild-type (WT). (C) Sequences of CRISPR-Cas9 induced indels from representative *tyr-*crispants. (Top) Mut1 and Mut4-7 sequences with deletions. (Below) Mut2 and 3 sequences with insertions. Targeted guide sites A (blue), B (pink), and C (cyan) are highlighted in wildtype reference sequences. *tyr-*crispants deletions/insertions are indicated in black and sequences matching wildtype in grey.

### Follicle train assignment

Follicles were ranked by size and by alternating left and right ovarian contributions to infer a probable timeline of egg lay for a given follicle size at time of injection. This method of follicle train assignment assumes 1) eggs present in the oviduct will be laid within one week, 2) follicles greater than 8-10 mm in 2 weeks, and 3) follicles less than or equal to 5mm in diameter will be laid no sooner than 3 weeks. Our reason for including these 3 underlying assumptions derives from the observation that females who had 1-2 eggs present in their oviduct, also had a follicle greater than 8-10 mm in diameter followed by a follicle between 3-5 mm in each ovary, suggesting at least a week interval between these sizes. As each lizard possesses a “leading” ovary and “lagging” ovary in follicle sizes, the leading ovary is given preferential ordering in train position. It is important to note that this method of ordering does not account for any potential loss of follicles accidently destroyed in the microinjection process and assumes that if such an event occurred, the follicle developmental timeline of that ovary is unaffected.

### Data availability

The data that support the findings of this study are available from the corresponding author upon request.

